# LL-37 and citrullinated-LL-37 enhance disparate oxylipins: LL-37-mediated chemokine response is dependent on COX-2 and the P2X_7_ receptor in human bronchial epithelial cells

**DOI:** 10.1101/2025.10.30.685679

**Authors:** Padmanie Ramotar, Mahadevappa Hemshekhar, Anthony Altieri, Anne M van der Does, Christopher Pascoe, Neeloffer Mookherjee

## Abstract

**Background:** During airway inflammation, chemokines, oxylipins (bioactive lipids) and cationic host defence peptides (CHDP) are enhanced in the lungs. However, the interplay of these molecules in the process of airway inflammation is not fully resolved. The human cathelicidin CHDP, LL-37, can enhance the expression of chemokines which is turn facilitates influx of leukocytes into the lungs. Moreover, LL-37 can get citrullinated during inflammation and the effect of this post-translational modification on LL-37-mediated immunomodulatory functions remains unclear. Therefore, in this study we aimed to define the impact of LL-37 and citrullinated-LL-37 (citLL-37) on oxylipins and its association with downstream chemokine production in human bronchial epithelial cells (HBEC), and its functional impact on leukocyte influx.

**Methods:** We used a lipidomics approach to identify oxylipins that are enhanced in response to LL-37 and citLL-37 in HBEC. We further examined the role of selected oxylipins in LL-37- and citLL-37-mediated chemokine production by ELISA, and related leukocyte migration using a transwell migration assay.

**Results:** We showed that LL-37, but not citLL-37, enhances oxylipins that are known to promote inflammation such as prostaglandins regulated by the cyclooxygenase (COX pathway). Although both LL-37 and citLL-37 upregulated COX-2, LL-37-mediated increase in COX-2 expression was significantly higher than that mediated by citLL-37. We showed that upregulation of COX-2 expression was dependent on the P2X_7_ purinergic receptor. Our mechanistic studies revealed that LL-37-mediated increase in chemokines, GROα, IL-8 and MIP-3α, was dependent on the COX-2 pathway. Our results also indicated that COX-2-induced PGE_2_ may act in an autocrine manner signaling through its EP receptors to facilitate LL-37-induced chemokine production. We functionally confirmed that factors secreted from HBEC in response to LL-37, but not citLL-37, promotes neutrophil migration which is COX-2 dependent.

**Conclusion:** The results of this study indicate that pro-inflammatory responses mediated by LL-37 is alleviated by citrullination of the peptide. These findings suggest that citrullination of LL-37 may be a post-translational regulatory mechanism to control inflammation. Overall, this study underscores the role of LL-37 in influencing the enhancement of bioactive lipids and metabolic pathways such as COX-2 and its link to the peptide-mediated immunomodulatory functions in the lungs.

## Background

Inflammation is a critical component of immune response that involves an intricate network of cellular and molecular events. During airway inflammation, human bronchial epithelial cells (HBEC) releases chemokines that facilitate recruitment of leukocytes such as neutrophils to the lungs. In addition, during airway inflammation immunomodulators such as oxylipins and cationic host defense peptides (CHDP) are also released in the lungs, which play a role in the modulation of inflammation (1–3). Although chemokines, oxylipins and CHDPs are all released in the lungs, the interplay of these immunomodulators remain largely unexplored in the process of airway inflammation. Thus, the focus of this study was to characterize oxylipins enhanced in response to the human CHDP LL-37 in HBEC, and its impact on LL-37-mediated chemokine production and neutrophil migration.

Oxylipins are bioactive lipids that can facilitate inflammation (1), generated from the oxidation of polyunsaturated acids such as arachidonic acid which is released from the lipid membrane by phospholipase A_2_ (4). Arachidonic acid serves as a substrate for three different metabolic pathways to produce oxylipins; the cyclooxygenase (COX) pathway which generates prostaglandins, the lipoxygenase (LOX) pathway which produces leukotrienes, and the cytochrome P450 monooxygenases pathway which produces epoxides (5). Oxylipins bind to specific G protein-coupled receptors (GPCRs) to mediate downstream signaling pathways related to immune response and inflammation (1). Prostaglandin E_2_ (PGE_2_) and the COX-2 pathway are the most well characterized mediators implicated in the process of inflammation, including in airway inflammation (6). Previous studies have demonstrated that influence of LL-37 on oxylipins in different cell types. For example, LL-37 was shown to enhance PGE_2_ in endothelial cells, fibroblasts and keratinocytes (7–9), and induce leukotriene production in eosinophils (10) and macrophages (11). Thus, to provide a more detailed insight into the impact of LL-37 on oxylipin production in airway inflammation, we used a lipidomics approach to identify oxylipins that are enhanced in response to LL-37 in HBEC.

A consideration in the biological function of LL-37 in airway inflammation is the aspect of post-translational modification of the peptide. Citrullination is a post-translational modification wherein the positively charged arginine residues of LL-37 are converted to neutral citrulline by peptidyl-arginine deiminases (12,13). Consequently both LL-37 and citrullinated forms of LL-37 are found in the lungs (12,13). While citrullination of LL-37 has been shown to impair its antimicrobial and antiviral properties (14,15), the effect of citrullination on the immunomodulatory functions of LL-37 remains largely elusive. We have recently shown that citrullination does not impair all immunomodulatory functions of LL-37, instead it selectively suppress the pro-inflammatory functions of LL-37 (16,17). Therefore, in this study we also compared the effect of LL-37 and citrullinated-LL-37 (citLL-37) on oxylipins and chemokine production, and functionally on neutrophil migration.

Overall, in this study we define oxylipins that are enhanced in response to LL-37 and citLL-37 in HBEC. We demonstrate that these peptides differentially upregulate COX-2 expression engaging the P2X_7_ purinergic receptor. Our findings indicate that citrullination of LL-37 selectively suppresses the ability of the peptide to induce pro-inflammatory oxylipins, COX-2 expression, and chemokines in HBEC. We functionally demonstrate that citrullination of LL-37 mitigates the ability of the peptide to facilitate neutrophil migration. The findings of this study indicate that citrullination may be a molecular switch to control LL-37-mediated inflammatory responses in the lungs. Notably, this study highlights the role of LL-37 in influencing metabolic pathways to facilitate the peptide’s immunomodulatory functions, thus defining a role for LL-37 in immunometabolism.

## Methods

### Reagents

The cationic peptides LL-37, citLL-37, and scrambled LL-37 (sLL-37) were synthesized and purchased from Innovagen AB (Lund, Sweden) and stored in −20°C until use. Peptides were reconstituted in endotoxin-free water (Fisher Scientific, Cat# SH3052901), aliquoted and stored in glass vials at −20°C for a maximum of 3 months. Each reconstituted peptide aliquot was thawed at room temperature (RT), sonicated for 30 seconds in a water bath and vortexed for 15 seconds before use. Human recombinant IL-8 (carrier-free) was purchased from R&D Systems (Oakville, ON, CA; Cat# 208-IL), aliquoted and stored at −80°C until use. The COX-2 inhibitor Rofecoxib was obtained from Selleckchem (Burlington, ON, CA; Cat# S3043), reconstituted in dimethyl sulfoxide (DMSO; Fisher Scientific) according to the manufacturer’s instructions. P2X_7_ inhibitor, KN62 (Cat# 13318) and PGE_2_ receptors (EP_1-4_) inhibitors, SC-19220 (EP_1_; Cat# 14060), PF-04418948 (EP_2_; Cat# 15016), L-798,106 (EP_3_; Cat# 11129) and MF498 (EP_4_; Cat# 15973), were purchased from Cayman Chemical (Ann Arbor, MI, USA) and reconstituted in DMSO (Fisher Scientific) according to the manufacturer’s instructions.

### Human bronchial epithelial cell cultures; cell line and human primary cells

The human bronchial epithelial cell line HBEC-3KT (CRL-4051™) was purchased from American Type Culture Collection, cultured in airway epithelial cell basal medium (ATCC® PCS-300-030™) supplemented with bronchial epithelial cell growth kit (ATCC® PCS-300-040™), according to the manufacturer’s instructions and as previously described by us (16). The cell culture medium was replaced with airway epithelial cell basal medium containing 6 mM L-glutamine, without the growth supplements, 24 hours (h) prior to stimulation with peptides in the presence or absence of various inhibitors. HBEC-3KT cells were stimulated with either LL-37, citLL-37, and sLL-37 (concentrations as indicated), mRNA was isolated after 4 h and tissue culture (TC) supernatants were collected after 24 h.

Human primary bronchial epithelial cells (PBEC) were isolated from macroscopically normal lung tissue obtained from patients undergoing resection surgery for lung cancer at the Leiden University Medical Center (LUMC), the Netherlands, as previously described by us (16). Lung tissue donors were enrolled in the biobank via a no-objection system for coded anonymous further use of such tissue (www.coreon.org). Samples from this Biobank were approved for research use by the institutional medical ethical committee (BB22.006/AB/ab). Since 01-09-2022, patients are enrolled in the biobank using written informed consent in accordance with local regulations from the LUMC biobank with approval by the institutional medical ethical committee (B20.042/KB/kb). Briefly, human PBECs were expanded in T75 TC flasks precoated with coating media (10 μg/mL fibronectin (Sigma), 30 μg/mL PureCol (Advanced Biomatrix, California, USA) and 10 μg/mL bovine serum albumin [(BSA; Sigma) in phosphate buffered saline (PBS)] in keratinocyte serum-free medium (KSFM; Gibco) supplemented with 25 μg/mL of bovine pituitary extract (BPE; Gibco), 0.2 ng/mL of epidermal growth factor (EGF; Life Technologies), 1:100 dilution of antibiotics Penicillin and Streptomycin (Lonza, Kingston, ON, CA) and 1 μM isoproterenol (Sigma-Aldrich, Oakville, ON, CA). Cells were expanded to approximately 80% confluency, followed by trypsinization with 0.3 mg/mL trypsin in PBS containing 1 mg/mL glucose (Gibco), 0.1 mg/mL EDTA (Gibco) and 1:100 dilution of Penicillin and Streptomycin. PBEC were seeded at a density of 1×10^4^ cells/mL in 12-well TC plates (Costar^®^) precoated with coating media, and cultured in a 1:1 mixture of basal bronchial epithelial cell medium (ScienCell, CA, USA) containing bronchial epithelial cell growth supplement (ScienCell), 1 nM of EC-23 (Tocris, UK), along with Dulbecco’s modified Eagle’s medium (Gibco) containing a 1:40 dilution of HEPES buffer (Invitrogen), and 1:100 dilution of Penicillin and Streptomycin. The cell culture medium was replaced every 48 h until, the medium was replaced without EGF, BPE and BSA 24 h prior to stimulation with either peptides or inhibitors as indicated.

### Cytotoxicity assay

Cytotoxicity was determined by examining TC supernatants for the release of the lactate dehydrogenase (LDH) enzyme using a colorimetric assay (Roche Diagnostic, Laval, QC, CA), according to the manufacturer’s instructions. Briefly, TC supernatants were centrifuged (250 x g for 5 minutes (min)) to obtain cell-free samples. TC supernatants (50 μL) were incubated with the LDH substrate mix (50 μL) for ~30 minutes in dark at RT, followed by colorimetric detection at 490 nm. Cytotoxicity was calculated relative to TC supernatants obtained from cells treated with 2% Triton X-100 (Sigma) as a measure for 100% cytotoxicity.

### Oxylipin profiling with mass spectrometry

HBEC-3KT cells were stimulated with either LL-37, citLL-37 or sLL-37 (0.25 μM each) as indicated for 24 h. TC supernatants were centrifuged at 250 x g for 5 min to obtain cell-free samples. Oxylipin abundance was measured in TC supernatants using LC-MS/MS as previously described by us (20). Briefly, oxylipins were extracted from the TC supernatants with solid phase extraction using Strata-X SPE columns (Phenomenex, CA, USA) preconditioned with methanol and pH 3 water and eluted with methanol. The extracted oxylipins were dried down under a gentle stream of nitrogen and resuspended in solvent A (Water – Acetonitrile – Acetic Acid [70:30:0.02; v/v/v]) & Solvent B (Acetonitrile – Isopropyl Alcohol [50:50; v/v]). Samples were separated using reverse phase HPLC using a Luna column (Luna 5u C18(2) 100A; 250 x 2.00 mm, Phenomenex) with gradient elution using Solvent A (Water – Acetonitrile – Acetic Acid [70:30:0.02; v/v/v]) and Solvent B (Acetonitrile – Isopropyl Alcohol [50:50; v/v]), on a Nexera-XR LC-20AD XR (Shimadzu) HPLC. The HPLC was coupled to a Qtrap 6500 (SCIEX) mass spectrometer equipped with a IonDrive Turbo V Electrospray Ion source. Polarity was set to negative mode, and responses were measured using selective reaction monitoring (SRM). Pairwise differential analysis was conducted on normalized log_2_ oxylipins values. Oxylipins enhanced by sLL-37 were removed from the data analysis. Welch’s t-test with a cut-off of *p*<0.05 was used to select oxylipins that were significantly enhanced by LL-37 and citLL-37. Fold changes were computed for LL-37 and citLL-37-mediated oxylipin abundance relative to unstimulated cells.

### ELISA

Cells were stimulated with either LL-37, citLL-37 or sLL-37 as indicated, and TC supernatants were collected after 24 h. TC supernatants were centrifuged (250 x g for 5 min) to obtain cell-free samples. The abundance of IL-8 (Cat# DY208), GROα (Cat# DY275) and MIP-3α (Cat# DY360) were examined in TC supernatants using ELISA kits obtained from R&D Systems, according to the manufacturer’s instructions.

### Quantitative Real-Time PCR

Cells were stimulated with either LL-37, citLL-37 or sLL-37 as indicated for 2, 4, 6 or 24 h. The cells were washed with cold PBS and cell lysate collected using lysis buffer (ThermoFisher Scientific; Cat# AC149320050) containing MagMAX™ Lysis/Binding Solution Concentrate (ThermoFisher Scientific; Cat# AM8500). Total RNA was isolated from the cell lysates using the MagMAX™ RNA isolation kit (ThermoFisher Scientific; Cat# AM1830) and the KingFisher Flex711 Automated Extraction & Purification System-GL (ThermoFisher Scientific), according to the manufacturer’s instructions. Total RNA was quantified using a NanoDrop 2000 Spectrophotometer (ThermoFisher Scientific). Superscript III First-Strand Synthesis SuperMix for qRT-PCR was used for first strand cDNA synthesis (ThermoFisher Scientific; Cat# 11752050), according to the manufacturer’s instructions. QuantiTect Primer Assays (Qiagen) for CXCL8 (GeneGlobe ID QT00000322), CXCL1 (GeneGlobe ID QT00199752), CCL20 (GeneGlobe ID QT00012971), PTGS1 (GeneGlobe ID QT00210280), PTGS2 (GeneGlobe ID QT00040586) and 18s (GeneGlobe ID QT00199367) used for qRT-PCR. Fold change for each mRNA target was calculated using the comparative ΔΔCt method after normalization with 18s RNA as the reference gene, followed by Log_2_ transformation.

### Neutrophil isolation and migration assay

Human blood was obtained from healthy volunteers with informed written consent, approved by the University of Manitoba’s Human Research Ethics Board (Protocol #: HS11105, H2010:259). Venous blood was collected in EDTA vacutainer tubes and neutrophils isolated using the EasySep™ Direct Human Neutrophil Isolation Kit (STEMCELL technologies, Vancouver, BC, Canada; Cat# 19666), according to the manufacturer’s instructions and described by us (18). Briefly, 25 mL of blood was gently mixed with the isolation cocktail and 50 µL of RapidSpheres™ provided in the kit, followed by incubation at RT for 5 minutes. The beads were washed, and neutrophils isolated using negative selection.

HBEC-3KT cells were stimulated with either LL-37, citLL-37 or sLL-37 (0.50 μM each), in the presence and absence of Rofecoxib (20 nM), and TC supernatants were collected after 24 h. TC supernatants (600 μL) were added to the bottom chamber of a 12-well, 12 mm Transwell^®^ plate with 12 (5.0 μM Pore Polyester membrane) inserts (Costar^®^). The plates were incubated at 37 °C in a 5 % of CO_2_ incubator for 30 minutes. Subsequently, human blood-derived neutrophils (6 x 10^5^ cells/ insert, 200 μL) were added to the upper chamber of the Transwell insert. As a positive control, human recombinant chemokine IL-8 (30 ng/mL) was added to the bottom chamber of one of the wells in the Transwell plates containing airway epithelial cells basal medium (containing 6 mM L-glutamine), as previously described by us (18). The Transwell plates were incubated for 2 h at 37 °C, followed by counting the number of migrated neutrophils in the bottom chamber using a Scepter 3.0 Handheld Automated Cell Counter (Millipore Ltd, ON, Canada).

### Statistical analysis

All statistical analyses were performed using GraphPad Prism (version 10.1.1; GraphPad Software). Briefly, paired baseline correction was done for all experiments by subtracting values obtained from unstimulated cells. An unpaired t-test was used for qRT-PCR statistical analysis. Two-Way ANOVA was used to compare protein levels of IL-8, GROα, and MIP-3α between different experimental conditions in experiments with pharmacological inhibitors, both in the presence and absence of peptide treatments (LL-37, citLL-37, and sLL-37). A *p*-value of *p*<0.05 was considered statistically significant. All statistical analyses are detailed in each of the figure legends.

## Results

### LL-37 and citLL-37 disparately enhances oxylipins in human bronchial epithelial cells

As LL-37 enhances PGE_2_ in fibroblasts, keratinocytes and endothelial cells (7–9), we examined oxylipin profile in HBECs in response to LL-37 and citLL-37 using a lipidomics approach as previously described by us (20). HBEC-3KT cells were stimulated with the peptides LL-37, citLL-37 or sLL-37 (0.25 µM each) and TC supernatant was collected after 24 h (n=3). Concentration of the peptides used was based on our previous study demonstrating that 0.25 µM of LL-37 and citLL-37 enhances specific chemokines and cytokines in HBEC (17). The scrambled peptide, sLL-37, was used as a paired negative peptide control, as we have previously shown that sLL-37 does not elicit the immunomodulatory functions of LL-37 (17,21). Thus, any oxylipin enhanced in response to sLL-37 was removed from the data analysis. Our results demonstrated that prostaglandins were enhanced in response to LL-37 (>2-fold compared to unstimulated cells). LL-37 specifically enhanced the abundance of PGE_2_, 11b PGE_2_, Dihomo PGE_2_, Dihomo PGF_2_α, PGA_2_ and PGE_1_ (Table 1), all products of the COX pathway (6,22,23). Whereas none of these prostaglandins were enhanced in response to citLL-37 (Table 1). LL-37 also significantly enhanced the abundance of 11-HETE (>2-fold), 15-HETE (>26-fold) and 15-HETrE (>2-fold compared to unstimulated cells), oxylipins produced by both the COX and LOX pathways (Table 1). Whereas citLL-37 only enhanced the abundance of 15-HETrE significantly (Table 1). In contrast, both LL-37 and citLL-37 significantly enhanced the abundance of 9,10 EpOME (>5-fold compared to unstimulated cells) and 12, 13 EpOME (>23-fold compared to unstimulated cells), oxylipins that are products of the cytochrome P450 pathway (24,25). However, abundance of both 9,10 EpOME and 12, 13 EpOME enhanced in response to citLL-37 were >50% lower than that elicited in response to LL-37 (Table 1).

**Table 1:** Disparate enhancement of oxylipins by LL-37 and citLL-37 in human bronchial epithelial cells.

| Pathways | Oxylipins | LL-37 |  | citLL-37 |  |
| --- | --- | --- | --- | --- | --- |
|  |  | Avg Log<sub>2</sub><br>Fold Change | *p*-value | Avg Log<sub>2</sub><br>Fold Change | *p*-value |
| COX Pathway | PGE<sub>2</sub> | 2.63 ± 1.19 | 0.03 | 1.28 ± 0.144 | ns |
|  | 11b PGE<sub>2</sub> | 2.44 ± 1.04 | 0.03 | 1.27 ± 0.162 | ns |
|  | Dihomo PGE<sub>2</sub> | 2.34 ± 0.813 | 0.02 | 1.21 ± 0.291 | ns |
|  | Dihomo<br>PGF<sub>2</sub>α | 1.90 ± 0.415 | 0.03 | 1.15 ± 0.178 | ns |
|  | PGE<sub>1</sub> | 2.19 ± 0.9 | 0.04 | 1.16 ± 0.102 | ns |
| COX/LOX<br>Pathway | 11-HETE* | 3.47 ± 1.04 | 0.02 | 5.50 ± 3.00 | ns |
|  | 15-HETE | 26.47 ± 24.69 | 0.02 | 2.90 ± 0.92 | ns |
|  | 15-HETrE | 2.49 ± 0.73 | 0.02 | 2.39 ± 0.490 | 0.03 |
| Cytochrome<br>P450 Pathway | 9,10 EpOMe* | 5.83 ± 2.27 | 0.005 | 2.49 ± 0.522 | 0.02 |
|  | 12,13 EpOMe* | 23.60 ± 13.74 | 0.01 | 9.25 ± 3.15 | 0.004 |

\*Can also be produced non-enzymatically

### LL-37 and citLL-37 differentially upregulates COX-2 mRNA via the P2X_7_ receptor

As LL-37 significantly enhanced prostaglandins (Table 1), and as COX enzymes are critical in the production of prostaglandins, we next examined the COX-1 and COX-2 mRNA abundance in response to LL-37 and citLL-37. However, before examining COX expressions we needed to optimize the concentration of LL-37 within physiologically representative range for further mechanistic studies. Concentration of LL-37 in the lungs has been shown to be between 0.25 to 1 µM in airway inflammation (26). We have previously shown that LL-37 (0.25 and 0.5 µM) enhances the levels of chemokines IL-8, GROα and MIP-3α in HBEC-3KT cells (17). Therefore, we examined chemokine production in response to LL-37 (0.25 and 0.5 µM) in HBEC-3KT cells. Both 0.25 and 0.5 µM of LL-37 significantly enhanced the levels of IL-8, GROα and MIP-3α in TC supernatants after 24 h, however the response was more than 50% higher following stimulation with 0.5 µM compared to 0.25 µM of LL-37 (Supplementary Figure 1). Based on these results, we selected the concentration of 0.5 µM for peptide stimulation for further studies.

HBEC-3KT cells were stimulated with LL-37, citLL-37 or sLL-37 (0.5 µM each) for 4 h, and the mRNA abundance of COX-1 and COX-2 were determined by qRT-PCR (n=5). COX-1 mRNA abundance was not significantly enhanced by either LL-37 or citLL-37 (Figure 1a). Although both LL-37 and citLL-37 significantly enhanced the mRNA abundance of COX-2 compared to unstimulated cells, LL-37-mediated enhancement of COX-2 was ~50% higher than that mediated by citLL-37 (Figure 1b). These results were consistent with the lipidomics data showing an increase in prostaglandins (between 2 and 3-fold increase compared to unstimulated cells) in response to LL-37, but not significantly with citLL-37 (Table 1). These results suggest that citrullination of LL-37 suppresses the ability of the peptide to enhance elements in the COX-2 pathway.

**Figure 1.**
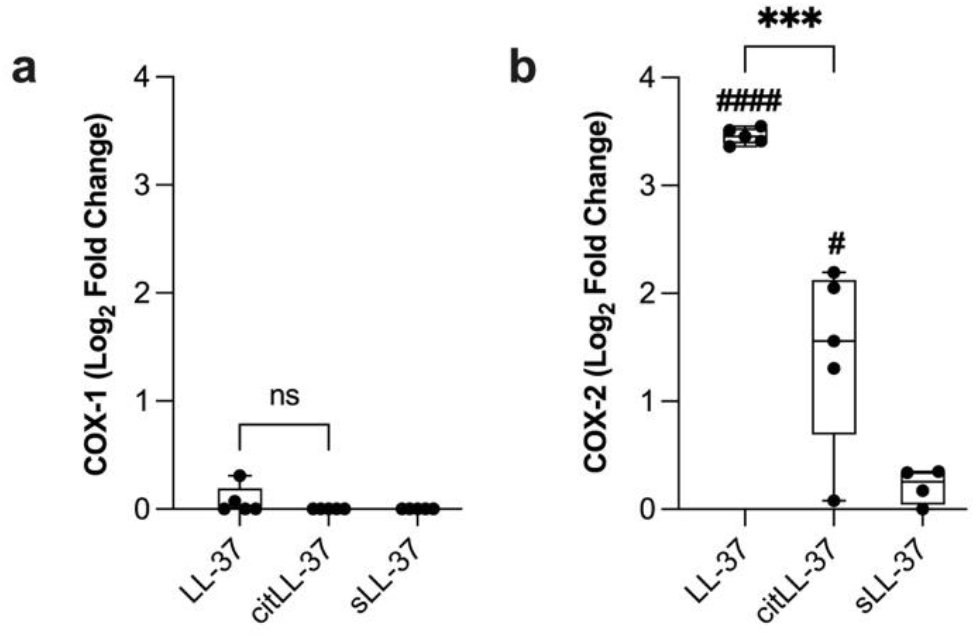
COX mRNA abundance in response to LL-37 and citLL-37. HBEC-3KT cells were stimulated with either LL-37, citLL-37 or sLL-37 (0.50 μM each). **(a)** COX-1 and **(b)** COX-2 mRNA abundance were examined in cell lysates using qRT-PCR after 4 h. Relative fold changes were calculated compared to unstimulated cells normalized to 1, using the ΔΔCt method after normalization with 18s RNA expression. Fold changes compared to unstimulated cells are shown after Log_2_ transformation. Each dot represents an independent experiment (n=5). Statistical significance was determined by paired t-test (# or * denotes *p*< 0.01, ### or *** denotes *p*<0.0005, and #### or **** denotes *p*<0.0001). # represents statistical significance compared to unstimulated cells

A previous study demonstrated that LL-37 engages the P2X_7_ purinergic receptor to induce COX-2 expression in human gingival fibroblasts (8). Therefore, we next examined the involvement of the P2X_7_ receptor in the enhancement of COX-2 mRNA, in response to LL-37 and citLL-37 in HBEC. HBEC-3KT cells were pre-treated with the P2X_7_ inhibitor, KN62 (10 nM) for 1 h, subsequently the cells were stimulated with either LL-37, citLL-37 or sLL-37 (0.5 µM each) for 4 h, and the mRNA abundance of COX-2 was determined by qRT-PCR. The inhibitor concentration was selected based on dose titration performed based on IC50 values provided by the manufacturer and existing literature (27) (Supplementary Figure 2). Cells were monitored for cytotoxicity by examining the release of LDH in TC supernatants. KN62 (10 nM) did not significantly increase cytotoxicity, in the presence or absence of the peptides, compared to unstimulated cells (data not shown). Inhibition of the P2X_7_ receptor significantly suppressed both LL-37 and citLL-37-mediated enhancement of COX-2 mRNA (Figure 2). These findings indicate that both LL-37 and citLL-37 engages the P2X_7_ receptor to enhance COX-2 mRNA in HBECs.

**Figure 2.**
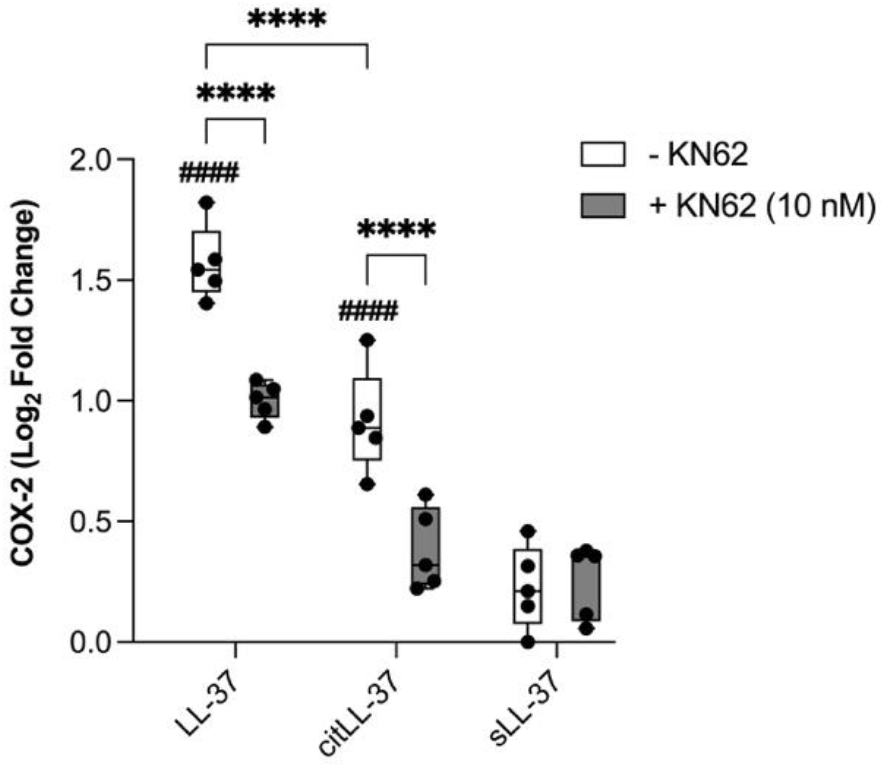
Inhibition of P2X_7_ receptor suppresses COX-2 mRNA. HBEC-3KT cells were pre-treated with KN62 (10 nM) for 1 h, followed by stimulation with either LL-37, citLL-37 or sLL-37 (0.50 μM each) for 4 h (n=5). Abundance of COX-2 mRNA was examined in cell lysates by qRT-PCR. Relative fold changes were calculated compared to unstimulated cells normalized to 1, using the ΔΔCt method after normalization with 18s RNA expression. Cells stimulated with KN62 were normalized to cells stimulated with inhibitor alone. Fold changes compared to unstimulated cells are shown after Log_2_ transformation. Each dot represents an independent experiment (n=5). Statistical significance was determined by Two-Way ANOVA (#### or **** denotes *p*<0.0001). # represents statistical significance compared to unstimulated cells.

### LL-37 and citLL-37-mediated chemokine response positively correlates with COX-2 expression

We showed that LL-37 and citLL-37 can upregulate the expression of COX-2, albeit with quantitative differences (Figure 1). The COX-2 pathway is associated with the production of chemokines (22,23), and we have previously shown that LL-37 enhances the levels of chemokines IL-8, GROα and MIP-3α in HBECs (17). Therefore, we examined the relationship of peptide-mediated upregulation of COX-2 and chemokines. HBEC-3KT cells were stimulated with LL-37, citLL-37 or sLL-37 (0.5 µM each) for 4 h, and the mRNA abundance of chemokines IL-8, GROα and MIP-3α, as well as COX-2, was determined by qRT-PCR. Similar to COX-2 (Figure 1b), LL-37 significantly enhanced mRNA abundance of IL-8, GROα and MIP-3α, and these chemokine responses were significantly less following stimulation with citLL-37 (Supplementary Figure 3).

Additionally, LL-37 and citLL-37-mediated increase in mRNA abundance of COX-2 showed a significant positive correlation with chemokine response, however, the magnitude of response by LL-37 was greatly reduced when the peptide was citrullinated (Figure 3).

**Figure 3.**
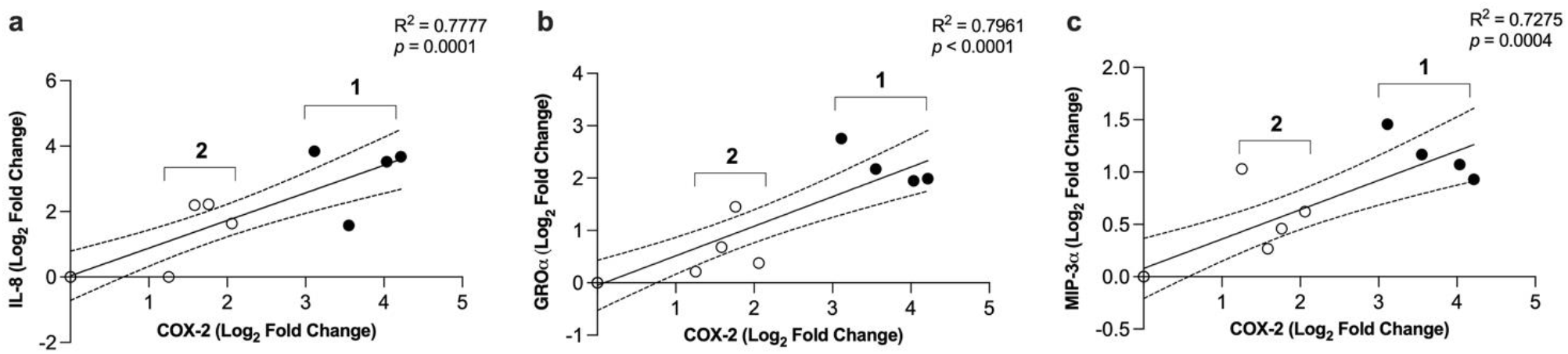
Correlations between LL-37 and citLL-37-mediated increase in the mRNA abundance of COX-2 and chemokines. HBEC-3KT cells were stimulated with either **(1)** LL-37 or **(2)** citLL-37 (0.50 μM each). mRNA abundance of COX-2, IL-8, GROα and MIP-3α were examined in cell lysates using qRT-PCR after 4 h. Relative fold changes were calculated compared to unstimulated cells normalized to 1, using the ΔΔCt method after normalization with 18s RNA expression. Pearson’s correlation analysis was performed to determine the correlation between COX-2 and **(a)** IL-8, **(b)** GROα and **(c)** MIP-3α mRNA abundance (fold changes compared to unstimulated cells). A *p*<0.05 was considered statistically significant

### LL-37-mediated chemokine production is dependent on COX-2

As there was a positive correlation between LL-37 and citLL-37-mediated upregulation of COX-2 and chemokines (Figure 3), we next examined the effect of COX-2 inhibition on peptide-mediated increase in IL-8, GROα and MIP-3α. HBEC-3KT cells were pre-treated with the COX-2 inhibitor Rofecoxib (20 nM) for 1 h. Concentration of the inhibitor was based on dose titration performed based on IC50 value provided by the manufacturer (Supplementary Figure 4). Subsequently, the cells were stimulated with LL-37, citLL-37 or sLL-37 (0.5 µM each) for 24 h and the protein abundance of IL-8, GROα and MIP-3α was examined in TC supernatants by ELISA. Cells were monitored for cytotoxicity by examining the release of LDH in TC supernatants under all conditions. The inhibitor did not increase cytotoxicity, in the presence or absence of the peptides, compared to unstimulated cells (data not shown). Presence of the COX-2 inhibitor significantly suppressed LL-37-mediated increase in the abundance of IL-8, GROα and MIP-3α proteins (Figure 4). In contrast, increase of GROα in response to citLL-37 was not significantly altered by inhibition of COX-2 (Figure 4). These results were also confirmed in human primary bronchial epithelial cells (PBEC) isolated from resected lung tissues. Like the results in HBEC-3KT cells (Figure 4), inhibition of COX-2 suppressed LL-37-mediated enhancement of chemokine production (Supplementary Figure 5). These results indicated that LL-37-mediated chemokine response is dependent on the COX-2 pathway. These results also aligned with the lipidomics data demonstrating enhancement of PGE_2_, an oxylipin downstream of COX-2, in response to LL-37 but not citLL-37 (Table 1).

**Figure 4.**
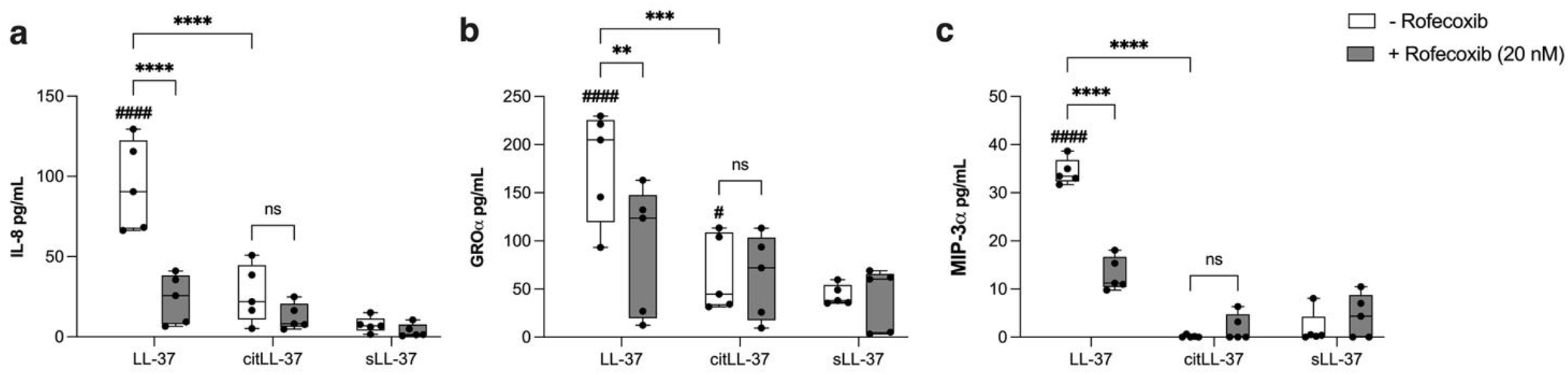
Inhibition of COX-2 suppresses LL-37-mediated chemokine production. HBEC-3KT cells were pre-treated with COX-2 inhibitor Rofecoxib (20 nM) for 1 h, followed by stimulation with either LL-37, citLL-37 or sLL-37 (0.50 μM each) for 24 h (n=5). TC supernatants were examined for the abundance of **(a)** IL-8, **(b)** GROα and **(c)** MIP-3α by ELISA. Chemokine levels shown in pg/mL after background subtraction of levels in unstimulated cells. Each dot represents an independent experiment (n=5). Statistical significance was measured using Two-Way ANOVA (# or * denotes *p*<0.05, ## or ** denotes *p*< 0.005, ### or *** denotes *p*< 0.0005, #### or **** denotes *p*< 0.0001). # represents statistical significance compared to unstimulated cells

### LL-37-mediated neutrophil migration is dependent on the COX-2 pathway

LL-37-induced chemokines IL-8 and GROα are both known to facilitate neutrophil recruitment, and LL-37 was previously shown to promote neutrophil migration (28). Similarly, COX-2-derived PGE_2_ can promote neutrophil migration (29). Our results demonstrated that LL-37-mediated IL-8 and GROα production were dependent on COX-2 (Figure 4). In addition, citLL-37 also modestly increased the abundance of GROα, albeit more than 50% less than that elicited in response to LL-37 (Figure 4). Therefore, we performed a functional assay to examine the impact LL-37 and citLL-37 on neutrophil migration, and its dependency on the COX-2 pathway. We examined the effect of a COX-2 inhibitor on neutrophil migration mediated by the secreted milieu of HBECs treated with the peptides. HBEC-3KT cells were pre-treated with the COX-2 inhibitor Rofecoxib (20 nM) for 1 h. Subsequently, the cells were stimulated with either LL-37, citLL-37 or sLL-37 (0.5 µM each) for 24 h. TC supernatants collected from these cells were used in the lower chamber of a Transwell plate for a neutrophil migration assay, as previously described by us (18). Neutrophils isolated from human blood were placed on the upper chamber of the Transwell plates, and the number of neutrophils in the lower chamber was counted after 3 h. TC supernatants obtained from HBEC-3KT cells treated with LL-37 significantly enhanced neutrophil migration, and this was significantly suppressed to baseline levels in the presence of the COX-2 inhibitor (Figure 5a).

**Figure 5.**
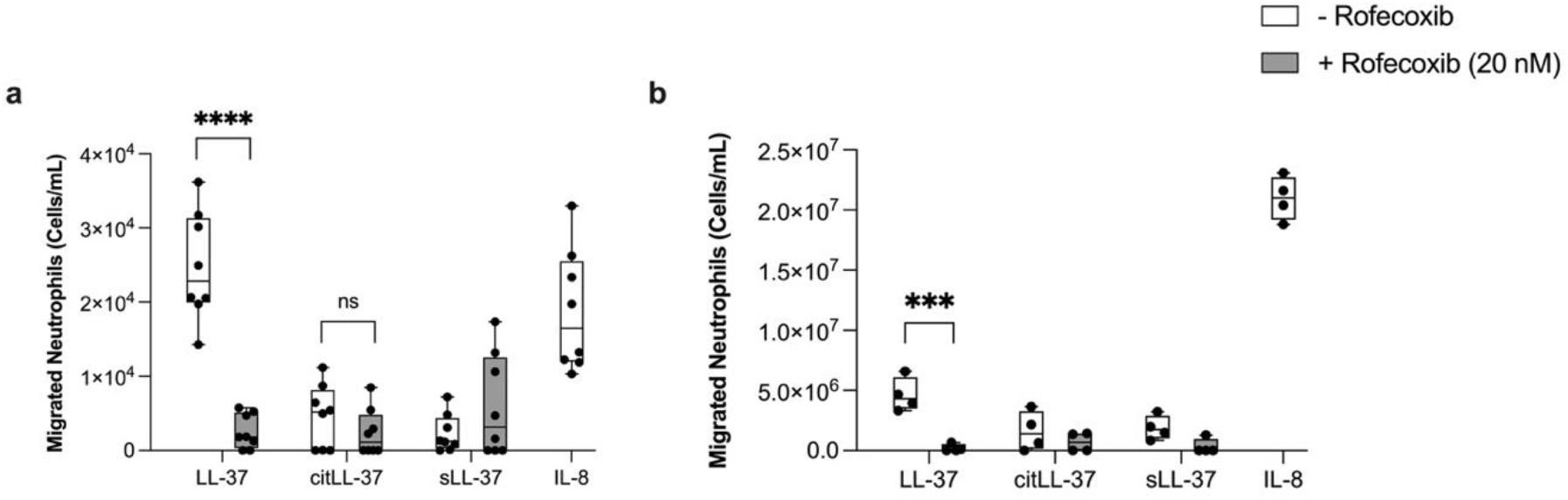
LL-37-mediated neutrophil migration is dependent on the COX-2 pathway. **(a)** HBEC-3KT cells and **(b)** human primary bronchial epithelial cells (PBEC) were pre-treated with the COX-2 inhibitor Rofecoxib (20 nM) for 1 h, followed by stimulation with either LL-37, citLL-37 or sLL-37 (0.50 μM each). TC supernatants collected after 24 h were used in the bottom chamber of a Transwell plates. IL-8 (30 ng/mL) was used in the bottom chamber of an independent well as a positive control for neutrophil migration. Neutrophils isolated from human blood, from two independent donors with two technical replicates each, were placed in the upper chamber of the Transwell plate. Neutrophils were counted in the bottom chamber of the Transwell plate after 3 h. Results shown are after subtracting background counts of neutrophils in the bottom chamber in wells with TC supernatants from unstimulated cells. Each dot represents an independent replicate. Statistical significance was measured using Two-Way ANOVA (\*\*\**p*<0.001, and \*\*\*\**p*<0.0001)

**Figure 6.**
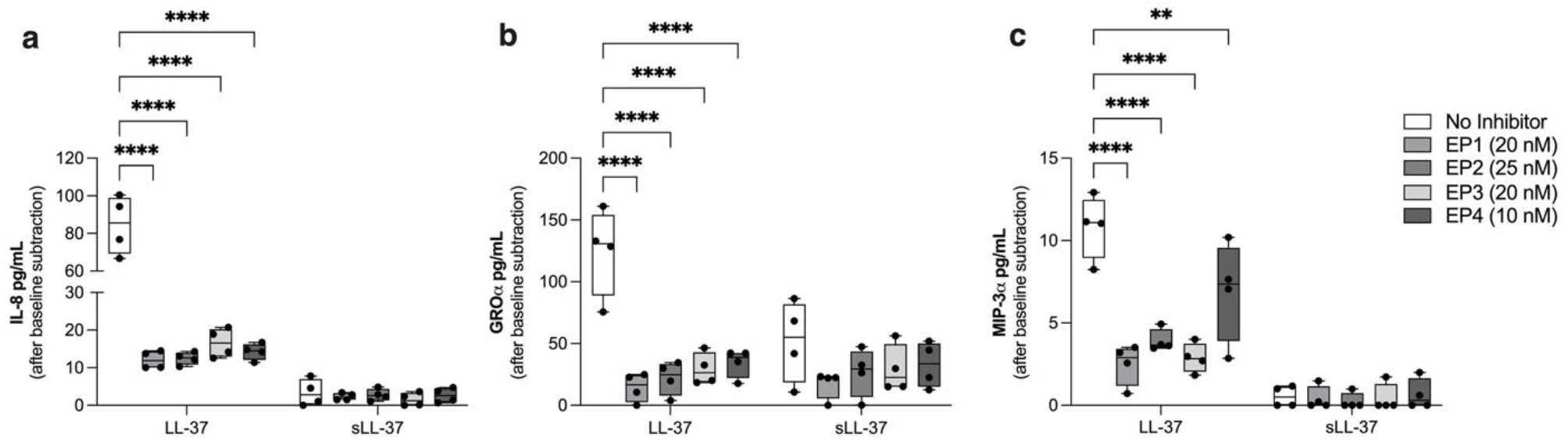
Inhibition of PGE_2_ receptors (EP_1-4_) suppresses LL-37-mediated enhancement of chemokines. HBEC-3KT cells were pre-treated with specific inhibitors for PGE_2_ receptors, EP_1_ (SC-19220; 20 nM), EP_2_ (PF-044EP2; 25 nM), EP_3_ (L-798,106; 20 nM), or EP_4_ (MF498; 10 nM), 1 h prior to stimulation with either LL-37or sLL-37 (0.50 μM). Tissue culture (TC) supernatants were collected after 24 h and the abundance of **(a)** IL-8, **(b)** GROα and **(c)** MIP-3α were measured by ELISA. Each dot represents an independent experiment (n=4). Results shown are after subtracting baseline values obtained from unstimulated cells in each independent experiment. Statistical significance was determined using Two-Way ANOVA (\*\**p*< 0.001, \*\*\*\**p*< 0.0001)

We further confirmed these results in primary cells using human PBEC isolated from lungs. PBECs were pre-treated with COX-2 inhibitor for 1 h, followed by stimulation of the cells with the peptides, as indicated above, for 24 h. TC supernatants obtained from PBECs under different conditions was used in the lower chamber of a Transwell plate for the neutrophil migration assay detailed above. Similar to the results in HBEC-3KT cells, TC supernatants obtained from human PBEC stimulated with LL-37 significantly enhanced neutrophil migration which was significantly inhibited in the presence of the COX-2 inhibitor (Figure 5b).

In contrast, TC supernatants obtained from both HBEC-3KT cell and human PBEC treated with citLL-37 did not promote neutrophil migration (Figure 5). These results indicated LL-37’s ability to facilitate neutrophil migration is dependent on the COX-2 pathway, and that citrullination of the peptide results in loss of function to facilitate neutrophil migration.

### Inhibition of PGE_2_ receptors suppresses LL-37-mediated enhancement of chemokines

Previous studies have shown that COX-2-mediated PGE_2_ can act in an autocrine manner via EP receptors to induce the production of chemokines, including IL-8 (30). We demonstrated that LL-37-mediated chemokine production, but not citLL-37, was dependent on the COX-2 pathway (Figure 4), and that LL-37 facilitates neutrophil migration (Figure 5). Therefore, we further examined whether LL-37-mediated chemokine production was dependent on the PGE_2_ receptors EP_1-4_. HBEC-3KT cells were treated with specific inhibitors for PGE_2_ receptors, EP_1_ (SC-19220; 20 nM), EP_2_ (PF-044EP2; 25 nM), EP_3_ (L-798,106; 20 nM), and EP_4_ (MF498; 10 nM), for 1 h. The inhibitor concentration was selected based on IC50 values provided by the manufacturer. In the presence or absence of the peptides, the inhibitors did not increase cytotoxicity compared to unstimulated cells (data not shown). Subsequently, the cells were stimulated with either LL-37 or sLL-37 (0.5 µM each) for 24 h, and the protein abundance of IL-8, GROα, and MIP-3α in TC supernatants was measured by ELISA. Inhibition of PGE_2_ receptors EP_1-4_ significantly suppressed LL-37-mediated increase of chemokines IL-8, GROα and MIP-3α. These results indicated that LL-37-mediated release of PGE_2_ (Table 1) may signal in an autocrine manner via the EP_1-4_ receptors to enhance chemokines production in HBECs.

## Discussion

The findings of this study provide an insight into differential production of oxylipins by LL-37 and the modified citrullinated peptide. Our findings show that LL-37 but not citLL-37 robustly enhances the production of prostaglandins in HBECs. We further demonstrate that LL-37 upregulates the expression of COX-2, which is significantly higher than that mediated by citLL-37. We also show that LL-37-mediated chemokine production and neutrophil migration is dependent on COX-2, a key enzyme within the metabolic pathway that produces prostaglandins. Overall, the findings of this study highlight a role for LL-37 in immunometabolism, linking bioactive lipids oxylipins produced by bronchial epithelial cells to the peptide’s functions in modulating chemokine responses and facilitating neutrophil migration, relevant to airway inflammation. Our findings also indicate that citrullination of LL-37 significantly suppresses the peptide’s ability to enhance COX-2, pro-inflammatory oxylipins and chemokines in HBECs, and mitigates LL-37-mediated neutrophil migration. These findings suggest that citrullination during airway inflammation may be a post-translational regulatory mechanism to control the pro-inflammatory functions of LL-37.

Both LL-37 and oxylipins function as immunomodulators that can either promote or suppress inflammation depending on the cellular milieu (31,32). Some previous studies have demonstrated an impact of LL-37 on oxylipins in different cell types. For example, LL-37 was shown to facilitate the release of PGE_2_ in endothelial cells (7), and upregulate the expression of COX-2 to promote PGE_2_ release in keratinocytes and gingival fibroblasts (8,9). These studies are corroborated by our findings demonstrating that LL-37 enhances COX-2 expression and PGE_2_ abundance in HBECs. A possible mechanism may be that LL-37 enhances oxylipin production through its involvement in lipid peroxidation. Oxylipins are products of polyunsaturated fatty acids (PUFA) oxidation, a complex process where PUFAs undergo reactions with oxygen to form new compounds (31).

Reactive oxygen species (ROS) are critical in initiating lipid peroxidation (33). LL-37 has been shown to facilitate increase in ROS production in neutrophils and macrophages (34). Thus, it is possible that LL-37 enhances ROS levels in the lungs and promotes oxidation of PUFAs to generate oxylipins, which warrants further investigation. Additionally, our lipidomics data demonstrate differential enhancement of other oxylipins such as those in the LOX and Cytochrome P450 pathways by LL-37 and citLL-37. While previous studies have shown that LL-37 can induce the release of leukotrienes (35), to our knowledge, this is the first study to show that LL-37 can enhance products from the LOX pathway, such as 11-HETE, 15-HETE, and 15-HETrE. To our knowledge, we also demonstrate for the first time that LL-37 enhances 9,10- and 12,13 EpOME, products of the CYP pathway. Although here we focus on the COX pathway, which regulates prostaglandins, involvement of the LOX and Cytochrome P450 pathways in mechanisms related to LL-37- and citLL-37-mediated immunomodulatory functions warrants further investigation, which is beyond the scope of this study.

In this study, we showed a direct association of COX-2 with LL-37-mediated enhancement of chemokines such as GROα and IL-8 that can facilitate neutrophilia. We confirmed this functionally by demonstrating that LL-37-mediated neutrophil migration is dependent on the COX-2 pathway. Thus, our findings clearly indicate that the pro-inflammatory functions of LL-37 to facilitate airway inflammation, in particular neutrophilic airway inflammation, may be dependent on the COX-2 pathway. Here, we also showed that the ability of LL-37 to enhance COX-2 is dependent on the P2X_7_ purinergic receptor in HBEC. This is consistent with a previous study demonstrating that LL-37 enhances COX-2 and PGE_2_ via the P2X_7_ receptor in gingival fibroblasts (8). However, we have previously shown that LL-37-mediated increase in chemokine production including GROα is dependent on GPCRs in macrophages (21). Interestingly, the biological functions of PGE_2_ is mediated by four different GPCRs subtypes EP_1-4_ (36). Previous studies have shown that COX-2-mediated PGE_2_ can act in an autocrine manner via EP receptors to induce the production of chemokines, including IL-8 (30). This is corroborated by our findings that inhibition of GPCRs EP_1-4_ receptors significantly suppresses the enhancement of chemokines in response to LL-37 in HBECs. Therefore, based on the results of this study we propose that LL-37 engages the P2X_7_ receptor on HBEC to enhance COX-2 expression facilitating the induction and release of PGE_2_, which subsequently acts in an autocrine manner via the PGE_2_ receptors (EP_1-4_), resulting in enhancement of chemokines such as IL-8 and GROα to facilitate neutrophil migration during airway inflammation (Figure 7).

**Figure 7.**
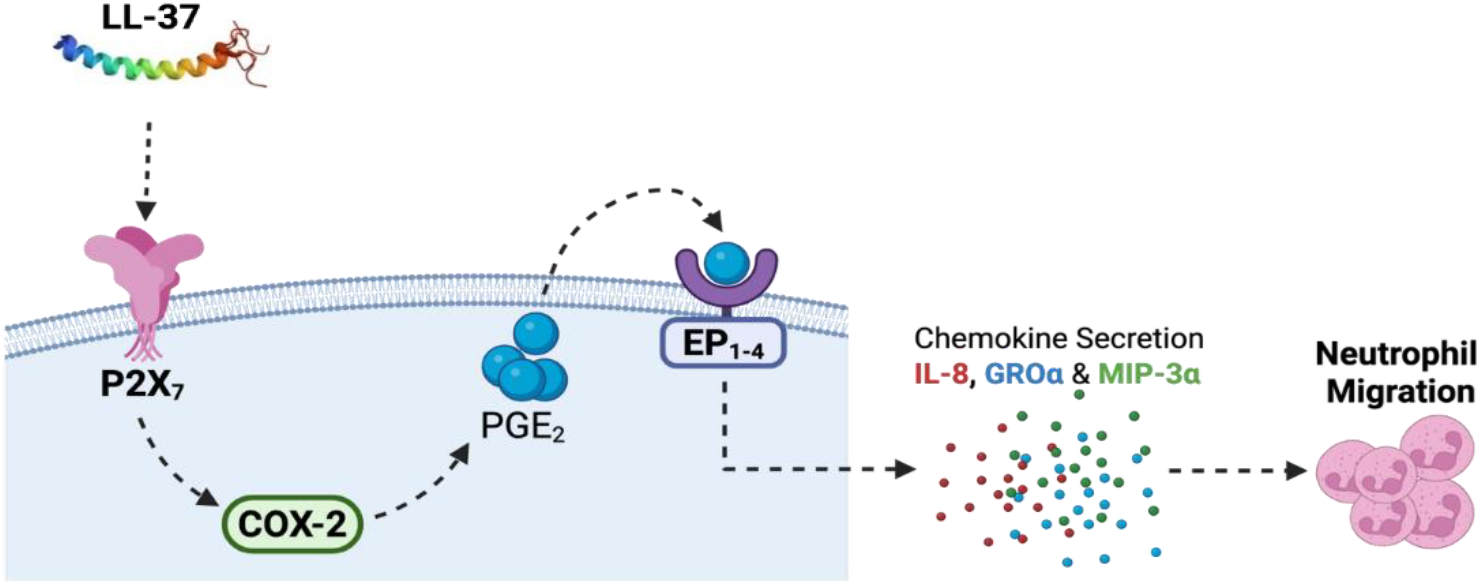
Proposed mechanism of LL-37-COX-2 axis for chemokine production and neutrophil migration in the lungs. LL-37 engages the P2X_7_ receptor in upregulating COX-2 expression which facilitates increase in the abundance of prostaglandin PGE_2_. Release of PGE_2_ may act in an autocrine manner through PGE_2_ receptors (EP_1-4_), resulting in enhanced production of chemokines including IL-8 and GROα, which facilitates neutrophil migration contributing to airway inflammation. However, citrullination of LL-37 dampens this pathway, potentially acting as a regulatory switch that limits the pro-inflammatory functions of LL-37 in the lungs (Figure created in BioRender.com).

As LL-37 becomes citrullinated during lung inflammation (12,14), it is important to compare the biological functions of LL-37 with that of citLL-37 to provide an insight into how the peptide activity may be altered in the lungs during airway inflammation. Our findings demonstrate that the ability of LL-37 to enhance pro-inflammatory oxylipins such as PGE_2_, chemokine production and facilitate neutrophil migration in the lungs is mitigated by citrullination. Previous studies have shown that citrullination of LL-37 results in impairment of LL-37’s antimicrobial functions (12,14,15). However, we have previously shown that citrullination of LL-37 does not abrogate all immunomodulatory functions of the peptide, instead selectively mitigates certain pro-inflammatory responses mediated by LL-37 (16,17). This is corroborated by the findings in this study demonstrating that although citLL-37 can modestly enhance COX-2 expression and GROα production, this is significantly less than that elicited by unmodified LL-37. We clearly show with a functional assay that citLL-37 is not able to promote neutrophil migration, which contrasts with LL-37. It is thus likely that citrullination of LL-37 during inflammation is a post-translational mechanism to regulate the peptide’s pro-inflammatory functions, potentially contributing to the resolution of the inflammatory process and restoring immune homeostasis.

Here, we show that the modest upregulation of COX-2 in response citLL-37 is also dependent on the P2X_7_ receptor. It is possible that the affinity of LL-37 to interact with P2X_7_ is reduced by citrullination, thereby impairing its ability to mediate downstream pro-inflammatory responses. It may also be that citLL-37 interacts with other receptors or protein partners which leads to the selective loss of pro-inflammatory responses of LL-37 (16,17). Studies defining the LL-37 interactome indicate that the peptide directly interacts with 16 different protein partners and indirectly with more than 1000 secondary effector proteins (37). How these interactions change following citrullination of LL-37 and consequent effects on the peptide’s immunomodulatory functions merits further systematic examination in the future.

A limitation of this study is that the findings are from *in vitro* experiments using undifferentiated HBECs, albeit using both cell line and human primary cells. A previous study showed that bronchial epithelial cells following mucociliary differentiation using air-liquid interface (ALI) cultures are more efficient in metabolic conversion of substrates in the generation of oxylipins (38). Nevertheless, the results of this study provide the foundation for future studies to systematically confirm the proposed mechanism (Figure 7) using ALI cultures and *in vivo* models of airway inflammation. Also, in this study we have used citLL-37 where all five arginine residues were modified to citrulline. However, under physiological conditions different forms of citrullinated LL-37 are detected in the lungs, and altered biological activity of LL-37 seem to be associated with the number of arginine residues of the peptide that are citrullinated (14). Therefore, the impact of citLL-37 on the peptide’s immunomodulatory functions and oxylipin production in the lungs may be quantitatively different depending on the number of arginine residues that are citrullinated, which also needs to be further examined.

## Conclusion

In this study, we have defined oxylipins that are enhanced in response to LL-37 and citLL-37 in human bronchial epithelial cells. We demonstrate that LL-37 enhances specific chemokines in HBECs, and that the resulting secreted milieu promotes neutrophil migration, via a COX-2-dependent mechanism. To our knowledge, this is the first study to link bioactive lipids and metabolic enzymes such as COX-2 to immunomodulatory functions of LL-37 in bronchial epithelial cells. This study underscores a role for LL-37 in immunometabolism during lung inflammation.

## List of abbreviations

(CHDP): Cationic host defence peptides
(citLL-37): citrullinated-LL-37
(COX): cyclooxygenase
(GPCRs): G protein-coupled receptors
(HBEC): human bronchial epithelial cells
(LDH): actate dehydrogenase
(LOX): lipoxygenase
(PGE_2_): Prostaglandin E_2_
(TC): tissue culture

## DECLARATIONS

### Ethics approval and consent to participate

Venous blood was obtained from healthy volunteers with informed written consent, approved by the University of Manitoba’s Human Research Ethics Board (Protocol #: HS11105, H2010:259). Human primary bronchial epithelial cells (PBEC) were isolated from macroscopically normal lung tissue obtained from patients undergoing resection surgery for lung cancer at the Leiden University Medical Center (LUMC), the Netherlands. Lung tissue donors were enrolled in the biobank via a no-objection system for coded anonymous further use of such tissue (www.coreon.org) and samples were approved for research use by the institutional medical ethical committee (BB22.006/AB/ab). Since 01-09-2022, patients are enrolled in the biobank using written informed consent in accordance with local regulations from the LUMC biobank with approval by the institutional medical ethical committee (B20.042/KB/kb).

### Consent for publication

Not applicable

### Availability of data and materials

All data generated and analyzed during this study are included in the published article and supplementary information.

### Authors’ contributions

PR and NM conceived and designed the study. PR performed majority of the experiments, analyzed the data, and wrote the manuscript. MH contributed to the optimization of some of the methods, provided intellectual input in study design and performed some of the experiments with primary cells. CP provided significant intellectual input for the study, performed the lipidomics sample processing and data analysis. AA performed initial optimization of peptides and chemokine endpoints in human bronchial epithelial cells. AMD provided the human primary bronchial epithelial cells and provided methodological guidance for primary cell cultures. NM obtained funding and resources for this study, provided overall supervision, and extensively edited the manuscript. All the authors reviewed the manuscript.

## Acknowledgments

We gratefully acknowledge Dr. Harold Aukema at the Canadian Centre for Agri-Food Research in Health and Medicine, for oxylipin profiling with mass spectrometry.

