## Supplemental Information for "LL-37 and citrullinated-LL-37 enhance disparate oxylipins: LL-37-mediated chemokine response is dependent on COX-2 and the P2X_7_ receptor in human bronchial epithelial cells"

### SUPPLEMENTARY INFORMATION

**Supplementary Figure 1** ***Concentration optimization of LL-37 for chemokine responses:*** HBEC-3KT were stimulated with LL-37 or sLL-37 (0.25 and 0.50 μM). Tissue culture (TC) supernatants were collected after 24 h. The abundance of **(a)** IL-8, **(b)** GROα, and **(c)** MIP-3α proteins were measured in TC supernatants by ELISA. Each dot represents an independent experiment (n=5). Statistical analysis was performed using Two-Way ANOVA (**$p< 0.001$ and ****$p< 0.0001$)

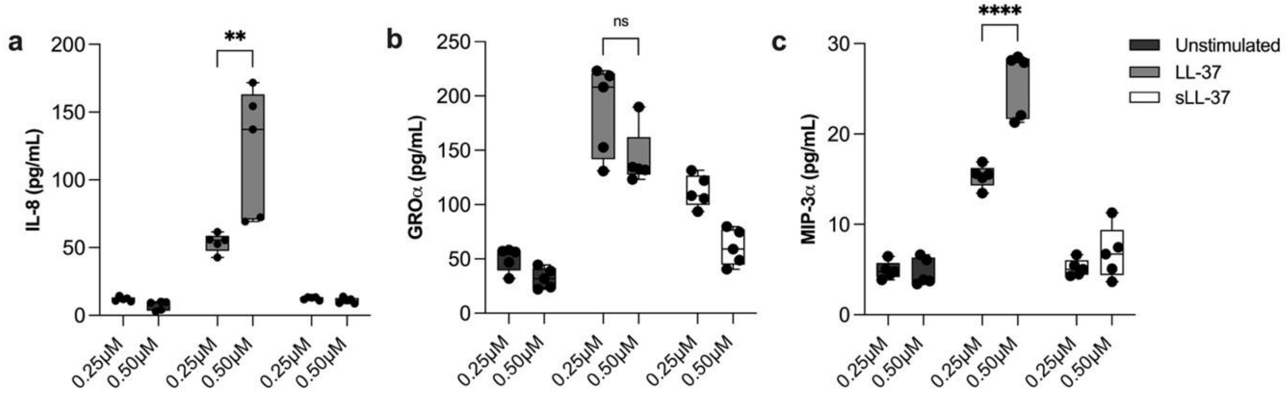


**Supplementary Figure 2** ***Optimization of concentration of P2X₇ inhibitor KN62.*** HBEC-3KT cells were pre-treated with the $P2X_7$ inhibitor KN62 (10 nM, 20 nM and 40 nM) for 1 h. Subsequently the cells were stimulated with peptides either LL-37, citLL-37 or sLL-37 (0.50 μM). Tissue culture (TC) supernatants were collected after 24 h. Protein abundance of **(a)** IL-8, **(b)** GROα and **(c)** MIP-3α were measured in TC supernatants by ELISA. Each dot represents an independent experiment (n=4). Statistical significance was determined using Two-Way ANOVA (***$p<0.0005$ and **** $p<0.0001$)

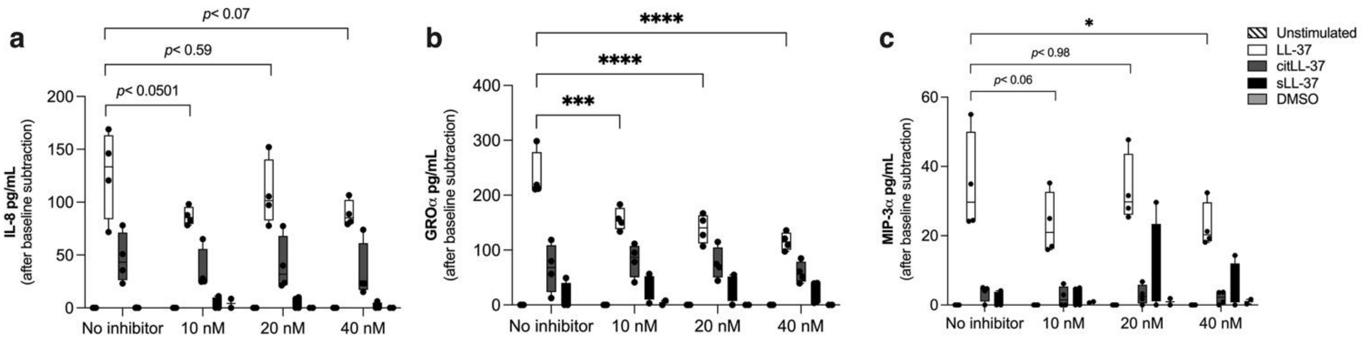

**Supplementary Figure 3** ***LL-37 and citLL-37-mediated enhancement of chemokine transcripts.*** HBEC-3KT cells were stimulated with either LL-37, citLL-37, or sLL-37 (0.50 μM). The mRNA abundance of **(a)** IL-8, **(b)** GROα and **(c)** MIP-3α was examined using qRT-PCR after 4 h. Relative fold changes were calculated compared to unstimulated cells normalized to 1, using the ΔΔCt method after normalization with 18s RNA expression. Results shown are with $Log_2$ transformation. Each dot represents an independent experiment (n=4), and statistical significance was determined by One-Way ANOVA (## or ** denotes *p*< 0.001, ### or *** denotes *p*<0.0005, and #### or **** denotes *p*< 0.0001). # represents statistical significance compared to unstimulated cells

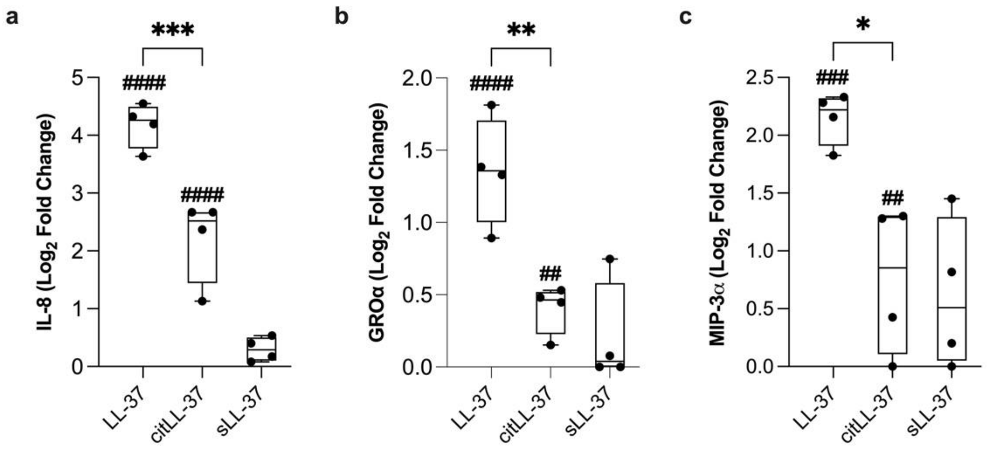


**Supplementary Figure 4** ***Optimization of COX-2 inhibitor concentrations.*** HBEC-3KT cells were pre-treated with COX-2 inhibitor Rofecoxib (10 nM, 20 nM or 40 nM) for 1 h. Subsequently the cells were stimulated with either LL-37, citLL-37 or sLL-37 (0.50 μM) for 24 h. Protein abundance of **(a)** IL-8, **(b)** GROα and **(c)** MIP-3α were measured in the tissue culture supernatants by ELISA. Each dot represents an independent experiment (n=2). Statistical significance was determined using Two-Way ANOVA (***p*<0.001, ****p*<0.0005 and **** *p*<0.0001)

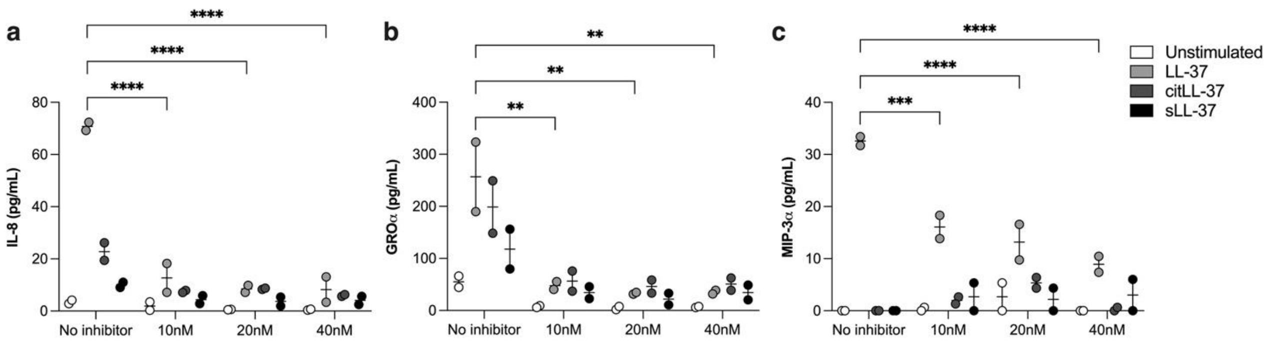


**Supplementary Figure 5** ***Inhibition of COX-2 suppresses LL-37-mediated chemokine production in human primary bronchial epithelial cells (PBEC)***. Human PBECs were treated with COX-2 inhibitor Rofecoxib (20 nM) for 1 h. Subsequently, the cells were stimulated with either LL-37, citLL-37 or sLL-37 (0.50 µM) for 24 h. The abundance of **(a)** IL-8, **(b)** GROα and **(c)** MIP-3α were examined in the tissue culture supernatants by ELISA. Results shown are after subtracting baseline values obtained in unstimulated cells from each condition, in each independent experiment. Each dot represents an independent experiment.

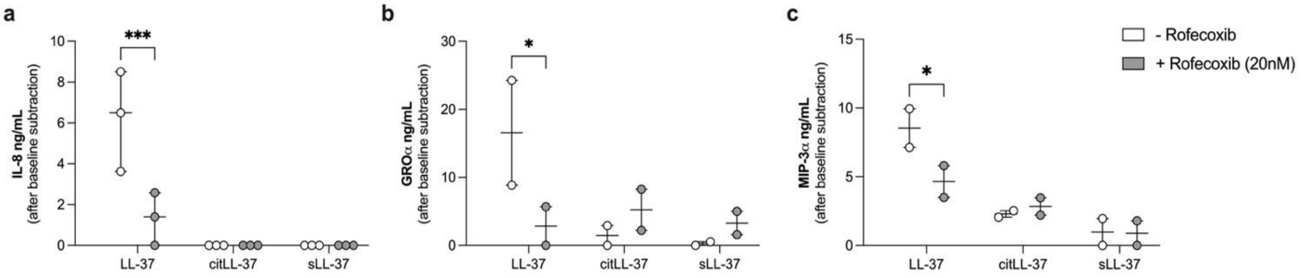
